# The effects of age on resting-state BOLD signal variability is explained by cardiovascular and cerebrovascular factors

**DOI:** 10.1101/836619

**Authors:** Kamen A. Tsvetanov, Richard N.A. Henson, P. Simon Jones, Henk-Jan Mutsaerts, Delia Fuhrmann, Lorraine K. Tyler, Cam-CAN, James B. Rowe

**Affiliations:** Department of Clinical Neurosciences, University of Cambridge, Cambridge, UK; Centre for Speech, Language and the Brain, Department of Psychology, University of Cambridge, Cambridge, UK; Medical Research Council Cognition and Brain Sciences Unit, Cambridge, UK; Department of Psychiatry, University of Cambridge, Cambridge, UK; Department of Radiology and nuclear medicine, Amsterdam University Medical Center, Amsterdam, The Netherlands

**Keywords:** ageing, functional magnetic resonance imaging (fMRI), individual differences, cerebral vascular reactivity

## Abstract

Accurate identification of brain function is necessary to understand neurocognitive ageing, and thereby promote health and well-being. Many studies of neurocognitive aging have investigated brain function with the blood-oxygen level-dependent (BOLD) signal measured by functional magnetic resonance imaging. However, the BOLD signal is a composite of neural and vascular signals, which are differentially affected by aging. It is therefore essential to distinguish the age effects on vascular *versus* neural function. The BOLD signal variability at rest (known as resting state fluctuation amplitude, RSFA), is a safe, scalable and robust means to calibrate vascular responsivity, as an alternative to breath-holding and hypercapnia. However, the use of RSFA for normalization of BOLD imaging assumes that age differences in RSFA reflecting only vascular factors, rather than age-related differences in neural function (activity) or neuronal loss (atrophy). Previous studies indicate that two vascular factors, cardiovascular health and cerebrovascular function, are insufficient when used alone to fully explain age-related differences in RSFA. It remains possible that their joint consideration is required to fully capture age differences in RSFA. We tested the hypothesis that RSFA no longer varies with age after adjusting for a combination of cardiovascular and cerebrovascular measures. We also tested the hypothesis that RSFA variation with age is not associated with atrophy. We used data from the population-based, lifespan Cam-CAN cohort. After controlling for cardiovascular and cerebrovascular estimates alone, the residual variance in RSFA across individuals was significantly associated with age. However, when controlling for both cardiovascular and cerebrovascular estimates, the variance in RSFA was no longer associated with age. Grey matter volumes did not explain age-differences in RSFA, after controlling for cardiovascular health. The results were consistent between voxel-level analysis and independent component analysis. Our findings indicate that cardiovascular and cerebrovascular signals are together sufficient predictors of age differences in RSFA. We suggest that RSFA can be used to separate vascular from neuronal factors, to characterise neurocognitive aging. We discuss the implications and make recommendations for the use of RSFA in the research of aging.

## 1. Introduction

The worldwide population is rapidly aging with an increasing number and proportion of older adults across the globe (Beard et al., 2016). Considering the cognitive decline and increasing burden of dementia in aging societies, there is a pressing need to understand the neurobiology of cognitive aging. This will inform efforts to maintain mental wellbeing into late life, allowing people to work and live independently for longer. Research in cognitive neuroscience of aging has used blood-oxygen level-dependent (BOLD) signal measured by functional magnetic resonance imaging (fMRI) as one of the standard ways to examine the neural mechanisms of cognition. However, the BOLD signal measures the activity of neurons indirectly through changes in regional blood flow, volume and oxygenation. This makes BOLD a complex convolution of neural and vascular signals, which are differentially affected by aging (Logothetis, 2008). Without careful correction for age differences in vascular health, differences in fMRI signals can be erroneously attributed to neuronal differences (Liu et al., 2013; Tsvetanov et al., 2015) and their behavioural relevance overstated (Geerligs et al., 2017; Geerligs and Tsvetanov, 2016; Tsvetanov et al., 2016).

It is possible to control for vascular differences in fMRI signal using additional baseline measures of cerebrovascular reactivity, including CO2-inhalation-induced hypercapnia (Liu et al., 2019), breath-hold-induced hypercapnia (Handwerker et al., 2007; Mayhew et al., 2010; Riecker et al., 2003; Thomason et al., 2007, 2005), hyperventilation-induced hypocapnia (Bright et al., 2009; Krainik et al., 2005), and cerebral blood flow (CBF) or venous oxygenation measures (Liau and Liu, 2009; Lu et al., 2010; Restom et al., 2007). However, such methods have not been widely used, in part to impracticalities in large-scale studies, and poor tolerance by older adults (for a review see Tsvetanov et al., 2020). Additionally, a hypercapnic challenge may not be neuronally neutral, given participants’ awareness of the aversive challenge, which may differ with age (Hall et al., 2011). Breath-hold compliance may also decrease with age (Jahanian et al., 2017). Such biases affect data quality and reliability measures (Magon et al., 2009), highlighting the advantage of non-invasive and “task-free” estimates of vascular components in the BOLD time series.

The BOLD signal variability in a resting state (“task-free”) is one such estimate and is also known as resting state fluctuation amplitudes (RSFA) (for a review see Tsvetanov et al., 2020). It has been proposed as a safe, scalable and robust cerebrovascular reactivity mapping technique (Golestani et al., 2016; Jahanian et al., 2014; Kannurpatti and Biswal, 2008; P. Liu et al., 2017). The use of RSFA as a normalization method for BOLD follows the assumption that age differences in RSFA reflect only vascular factors, rather than age-related differences in neural function or neuronal loss (atrophy). Fluctuations in the BOLD signal are associated with fluctuations in cardiac rhythm (Glover et al., 2007) that are independent of those associated with respiratory rate and depth (Chang et al., 2013, 2009), suggesting that RSFA may be susceptible to vascular signals of varying aetiologies, such as cardiovascular and cerebrovascular factors. Evidence in support of cardiovascular factors comes from Tsvetanov and colleagues (Tsvetanov et al., 2015, but also Makedonov et al., 2013; Viessmann et al., 2017, 2019; Theyers et al., 2018), who demonstrated that age-related differences in RSFA are mediated by cardiovascular health (as measured by pulseoximetry and electrocardiography, ECG), but not by neural function in terms of neural variability (as measured by magnetoencephalography, MEG). Evidence in support of cerebrovascular factors comes from Garrett et al. (2017) who found that “gold-standard” measures of cerebrovascular function (arterial spin labelling, ASL, and CO_2_ inhalation-induced hypercapnia) are associated with RSFA. Importantly, both studies reported age-related differences in RSFA that remain after adjusting for individual differences in either cardiovascular or cerebrovascular factors. However, neither study considered jointly cardiovascular and cerebrovascular factors, and it remains unclear whether the unexplained age-related differences in RSFA reflect joint contributions from cardiovascular and cerebrovascular factors, as in the case of BOLD signal fluctuations (Chang et al., 2013, 2009). Alternatively, the unexplained age differences in RSFA may reflect neuronal factors, such as atrophy (Grady and Garrett, 2013), even though variation in neuronal activity does not explain the effect of age on RSFA (Tsvetanov et al., 2015).

Cardiovascular, cerebrovascular and other physiological signals, but not neuronal signals, contribute to the age-related differences in RSFA, yet none of these non-neuronal measures on their own could fully account for the effects of age on RSFA. It is possible that various vascular signals contribute to different components of the age effects on RSFA (Tsvetanov et al., 2020). However, no study to date has tested whether the cardiovascular and cerebrovascular signals together fully capture the effects of age on RSFA – an assumption underlying the use of RSFA as a scaling method. In this study we sought to investigate the effects of age on RSFA by the simultaneous assessment of the independent and shared effects of cardiovascular, cerebrovascular and neuronal effects on age-related differences in RSFA. To this end, we used a set of cardiovascular, cerebrovascular and volumetric measures in a population-based study of healthy ageing (age 18-88, N > 250, www.cam-can.org). We hypothesized that age-related variation in RSFA are predicted by cardiovascular and cerebrovascular factors, but not grey matter volume, and therefore that the residuals in RSFA – after adjusting for these vascular factors – are not associated with age.

## 2. Methods

### 2.1. Participants

*Figure 1* illustrates the study design and image processing, using the Cambridge Centre Aging and Neuroscience dataset (Cam-CAN). Ethical approval was granted by Cambridgeshire 2 Research Ethics Committee. Participants gave written informed consent. A detailed description of exclusion criteria can be found in Shafto et al. (Shafto et al., 2014), including poor vision (below 20/50 on Snellen test; Snellen, 1862) or hearing (threshold 35dB at 1000Hz in both ears), ongoing or serious past drug abuse as assessed by the Drug Abuse Screening Test (DAST-20; Skinner, 1982), significant psychiatric disorder (e.g. schizophrenia, bipolar disorder, personality disorder) or neurological disease (e.g. stroke, epilepsy, traumatic brain injury). At an initial home assessment (Phase I), completed the Mini-Mental State Examination (MMSE > 25; Folstein et al., 1975) and Edinburgh Handedness Inventory (Oldfield, 1971). Participants attended MRI (T1-weighted, arterial spin labelling (ASL), FLAIR-based white matter hyperintensities, resting state EPI-BOLD and field-map images) and MEG (including resting state ECG-recording) on two occasions (Phase II and III) separated by approximately 1 year. We include here 226 full datasets of good quality, required for all analysis (e.g. T1-weighted, FLAIR, ASL, resting fMRI and ECG recordings, see below). Demographic characteristics of the sample are described in Table 1. Imaging data were acquired using a 3T Siemens TIM Trio.

**Table 1.** Participants’ demographic information, grouped by decile in accordance with the original design of the Cam-CAN cohort (Green et al., 2018; Shafto et al., 2014)

|  | Decile |  |  |  |  |  |  | Statistical tests* |  |
| --- | --- | --- | --- | --- | --- | --- | --- | --- | --- |
| | 1 | 2 | 3 | 4 | 5 | 6 | 7 | $\chi^2$ or F-test | P-value |
| **Age range [years]** | 18-27 | 28-37 | 38-47 | 48-57 | 58-67 | 68-77 | 78-90 |  |  |
| **Gender, n (% per decile)** |  |  |  |  |  |  |  | 0.15 | 0.989 |
| Men | 7 (46.7) | 19 (46.3) | 19 (50) | 19 (52.8) | 19 (50) | 17 (56.7) | 14 (50) |  |  |
| Women | 8 (53.3) | 22 (53.7) | 19 (50) | 17 (47.2) | 19 (50) | 13 (43.3) | 14 (50) |  |  |
| **Handedness\*\*** |  |  |  |  |  |  |  | 1.34 | 0.241 |
| Mean / SD | 91 / 12 | 85 / 42 | 86 / 27 | 93 / 11 | 79 / 48 | 97 / 5 | 91 / 30 |  |  |
| Range [Min/Max] | 65 / 100 | -65 / 100 | -56 / 100 | 58 / 100 | -78 / 100 | 86 / 100 | -56 / 100 |  |  |
| **Education, n (% per decile)** |  |  |  |  |  |  |  | 4.07 | <.001 |
| None | 0 (0) | 0 (0) | 0 (0) | 0 (0) | 0 (0) | 5 (16.7) | 1 (3.6) |  |  |
| GCSE/O-level | 2 (13.3) | 1 (2.4) | 6 (15.8) | 3 (8.3) | 4 (10.5) | 2 (6.7) | 4 (14.3) |  |  |
| A-level | 2 (13.3) | 2 (4.9) | 3 (7.9) | 12 (33.3) | 9 (23.7) | 9 (30) | 8 (28.6) |  |  |
| Degree | 11 (73.3) | 38 (92.7) | 29 (76.3) | 21 (58.3) | 25 (65.8) | 14 (46.7) | 15 (53.6) |  |  |
| **Mini-Mental State Exam** |  |  |  |  |  |  |  | 3.17 | 0.006 |
| Mean / SD | 29.5 / 0.9 | 29.6 / 0.7 | 29.1 / 1.2 | 29.2 / 0.9 | 29.1 / 1 | 28.7 / 1.3 | 28.8 / 1.3 |  |  |
| Range [Min/Max] | 27 / 30 | 27 / 30 | 26 / 30 | 26 / 30 | 27 / 30 | 26 / 30 | 25 / 30 |  |  |

\* Statistical test to indicate whether demographics vary between deciles
\*\* Higher scores indicate greater right-hand preference

**Figure 1.**
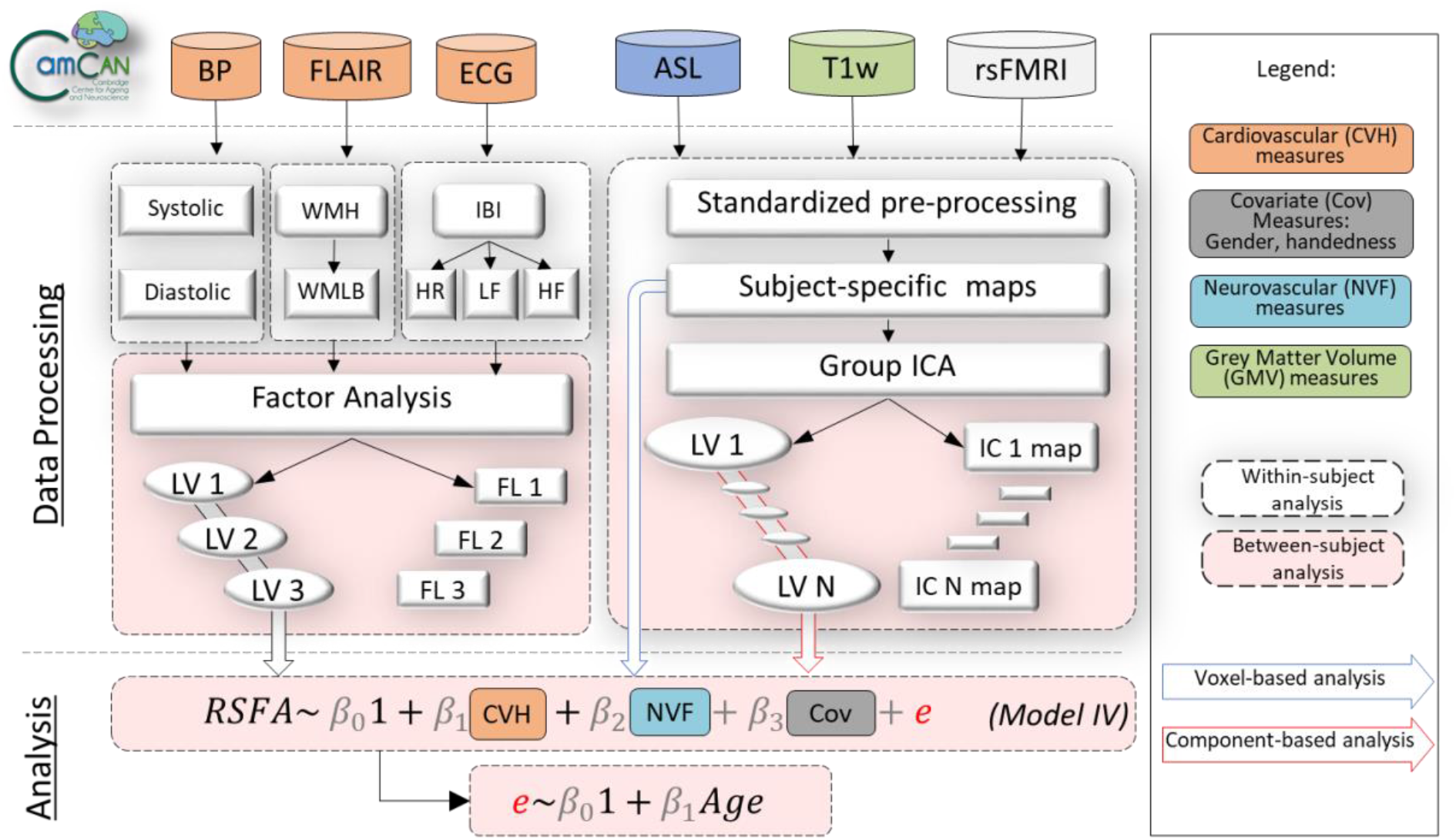
Visual representation of the analysis strategy in terms of data inclusion (above top dotted line), processing (below top dotted line) either at a within-subject level (white dotted-line rectangles) or between-subject level (peach-color dotted-line rectangles) and analysis (below second dotted line). Measures of cardiovascular health (CVH) included blood pressure (BP), heart rate variability (HRV) from electrocardiogram (ECG) recordings, white matter-matter hyperintensities (WMH) from fluid-attenuated inversion recovery (FLAIR) and BMI (not shown), all of which were submitted to factor analysis. Neurovascular function (NVF) estimates were based on cerebral blood flow from arterial spin labelling (ASL) acquisition. Grey matter volume (GMV) was estimated from a T1-weighted MRI acquisition. Resting state fluctuation amplitudes (RSFA) were estimated from resting-state fMRI BOLD acquisition. Regionally specific measures (RSFA, CBF and GMV) were submitted to multiple linear regression either on a voxel-level or on a component-level using outputs from group ICA. ICA – independent component analysis; LV – latent variable; LST – lesion-segmentation tool; PCA – principal component analysis; rsFMRI – resting state fMRI; TLV – total lesion volume; WMLB – white-matter lesion burden;

### 2.2. T1w image acquisition and processing

A 3D-structural MRI was acquired for each participant using T1-weighted Magnetization-Prepared Rapid Gradient-Echo (MPRAGE) sequence with Generalized Autocalibrating Partially Parallel Acquisition (GRAPPA) acceleration factor 2; Repetition Time (TR) = 2250ms; Echo Time (TE) = 2.99ms; Inversion Time (TI) = 900ms; flip angle α = 9°; field of view (FOV) = 256mm × 240mm × 192mm; resolution = 1mm isotropic) with acquisition time of 4 minutes and 32 seconds.

All image processing was done using Automatic Analysis (AA 4.0; Cusack et al., 2014; https://github.com/automaticanalysis/automaticanalysis) implemented in Matlab (Mathworks, https://uk.mathworks.com/). The results here come from Release004 of the CamCAN pipelines. Each particpant’s T1 image was coregistered to the MNI template in SPM12 (http://www.fil.ion.ucl.ac.uk/spm; Friston et al., 2007), and the T2 image was then coregistered to the T1 image using a rigid-body transformation. The coregistered T1 and T2 images underwent multi-channel segmentation (SPM12 Segment; Ashburner and Friston, 2005) to extract probabilistic maps of 6 tissue classes: GM, WM, cerebrospinal fluid (CSF), bone, soft tissue, and background. The native-space GM and WM segmentations were used for diffeomorphic registration (DARTEL; Ashburner, 2007) to create whole group template images (Taylor et al., 2015). The group template was normalised to the MNI space using 12-parameter affine transformation.

### 2.3. fMRI image acquisition and processing

RSFA was estimated from resting state Echo-Planar Imaging (EPI) of 261 volumes acquired with 32 slices (sequential descending order), slice thickness of 3.7 mm with a slice gap of 20% for whole brain coverage (TR = 1970ms; TE = 30ms; flip angle α = 78°; FOV = 192mm × 192mm; resolution = 3mm × 3mm × 4.44mm) during 8 minutes and 40 seconds. Participants were instructed to lay still with their eyes closed. The initial six volumes were discarded to allow for T1 equilibration. We quantified participant motion using the root mean square volume-to-volume displacement as per Jenkinson et al (2002). The rs-fMRI data were further pre-processed by wavelet despiking (see below).

The EPI data were unwarped (using field-map images) to compensate for magnetic field inhomogeneities, realigned to correct for motion, and slice-time corrected to the middle slice. The normalisation parameters from the T1 image processing were then applied to warp functional images into MNI space. We applied data-driven wavelet-despiking to minimise motion artefacts (Patel et al., 2014). We observed a high association between the amount of outlying wavelet coefficient and head motion across subjects (*r* = .739, *p* < .001), demonstrating that it captured a large amount of motion artefacts in the data. Spatially normalised images were smoothed with a 12 mm FWHM Gaussian kernel. A general linear model (GLM) of the time-course of each voxel was used to further reduce the effects of noise confounds (Geerligs et al., 2017), with linear trends and expansions of realignment parameters, plus average signal in WM and CSF, their derivative and quadratic regressors (Satterthwaite et al., 2013). The WM and CSF signal was created by using the average across all voxels with corresponding tissue probability larger than 0.7 in associated tissue probability maps available in SPM12. A band-pass filter (0.0078-0.1 Hz) was implemented by including a discrete cosine transform set in the GLM, ensuring that nuisance regression and filtering were performed simultaneously (Hallquist et al., 2013; Lindquist et al., 2019). Finally, we calculated subject specific maps of RSFA based on the normalized standard deviation across time for processed resting state fMRI time series data.

### 2.4. ASL image acquisition and processing

To assess resting cerebral blood flow, we used pulsed arterial spin labelling (PASL, PICORE-Q2T-PASL in axial direction, 2500ms repetition time, 13ms echo time, bandwidth 2232 Hz/Px, 256 × 256 mm^2^ field of view, imaging matrix 64×64, ten slices, 8 mm slice thickness, flip angle 90°, 700 ms inversion time (TI) 1, TI2 = 1800 ms, 1600 ms saturation stop time). The imaging volume was positioned to maintain maximal brain coverage with a 20.9 mm gap between the imaging volume and a labelling slab with 100mm thickness. There were 90 repetitions giving 45 control-tag pairs (duration 3’52”). In addition, a single-shot EPI (M0) equilibrium magnetization scan was acquired. Pulsed arterial spin labelling time series were converted to cerebral blood flow (CBF) maps using ExploreASL toolbox (Mutsaerts et al., 2018). Following rigid-body alignment, the spatial normalised images were smoothed with a 12 mm FWHM Gaussian kernel.

### 2.5. Cardiovascular measures

#### 2.5.1. Physiological recordings

Cardiac activity data were acquired using bipolar ECG while acquiring the MEG data, and processed using PhysioNet Cardiovascular Signal Toolbox (Goldberger et al., 2000; Vest et al., 2018) in Matlab (MATLAB 2017b, The MathWorks Inc, Natick, MA). To address non-stationarity in ECG recordings, mean heart rate (HR) and hearth rate variability (HRV) summary measures were based on the median across multiple sliding 5-min windows in 30-second steps across the entire eyes-closed, resting-state acquisition, 8.5 minutes. Estimation of mean heart rate (HR) was based on the mean number of successive N-N (normal-to-normal) intervals within each 60-second interval during each 5-minute period recording. To estimate the HRV, we used the frequency-domain information of normal-to-normal (NN) intervals, which provides a measure of low- and high-frequency components of the HRV (unlike time-domain alternatives e.g. the root mean squared difference of successive intervals (RMSSD), which pertain mainly to high-frequency dynamics of HRV, (Malik et al., 1996). We calculated low-frequency (0.05 – 0.15 Hz; LF-HRV) and high-frequency (0.15-0.4 Hz; HF-HRV) power. Segments classified as atrial fibrillation were excluded from further analysis, and any participant with >50% atrial fibrillation was excluded.

#### 2.5.2. White matter hyperintensities (WMH)

Estimates of white matter lesion burden in our sample have been reported previously (Fuhrmann et al., 2017). In summary, white matter lesion was estimated using the lesion growth algorithm in the LST toolbox for SPM (Schmidt et al., 2012) with *κ* of 0.7.

#### 2.5.3. Other risk factors of cardiovascular health: blood pressure and body mass index

Systolic and diastolic blood pressure were measured at rest, seated, using an automated sphygmomanometer (A&D Medical Digital Blood Pressure Monitor, UA-774). The average of three measurements was used. BMI was calculated as weight (kg) / height (m)^2^, using portable scales (Seca 875).

### 2.6. Data reduction

Datasets of interest stemmed from a wide range of modalities (RSFA, ASL, T1-weghted, FLAIR and ECG measures). To make these datasets tractable, we analysed a set of summary measures for each of the modality (also known as features or components) as illustrated in Figure 1. This had two advantages. First, it reduced the number of statistical comparisons. Second, it separated spatially overlapping sources of signal with different aetiologies within a modality (Xu et al., 2013), e.g. cardiovascular *versus* cerebrovascular signals, which may vary across individuals and brain region in RSFA (Tsvetanov et al., 2015) and ASL data (Mutsaerts et al., 2017). We used independent component analysis (ICA) across participants to derive spatial patterns of each imaging modality across voxels. As a proxy of vascular health, we used exploratory factory analysis to derive a latent variables from a set of measures related to cardiac function derived from the resting heart rate signal and other risk factors (Varadhan et al., 2009; Wardlaw et al., 2014).

#### 2.6.1. Indices of RSFA, T1 and CBF maps using Independent Component Analysis

Group ICA was implemented on RSFA, GMV and CBF maps separately. For each modality, data were decomposed to a set of spatially independent sources using the Source Based Morphometry toolbox (Xu et al., 2009) in the Group ICA for fMRI Toolbox (GIFT; http://mialab.mrn.org/software/gift).

In brief, the fastICA algorithm was applied after the optimal number of sources explaining the variance in the data was identified using PCA with Minimum Description Length (MDL) criterion (Hui et al., 2011; Li et al., 2007; Rissanen, 1978). By combining the PCA and ICA, one can decompose an n-by-m matrix of participants-by-voxels into a source matrix that maps independent components (ICs) to voxels (here referred to as “IC maps”), and a mixing matrix that maps ICs to participants. The mixing matrix indicates the degree to which a participant expresses a defined IC. The loading values in the mixing matrix were scaled to standardized values (Z-scores) and used for between-participant analysis of summary measures from other modalities. The maximum number of available datasets within each modality was used, recognising that ICA decomposition accurately represents individual variation despite different group sizes while maximizing statistical power (Calhoun et al., 2008; Erhardt et al., 2011).

#### 2.6.2. Indices of vascular health using Exploratory Factor Analysis

As a vascular health index, we sought a summary measure that characterized the complexity of cardiovascular signal (Varadhan et al., 2009; Wardlaw et al., 2014). We used factor analysis on the mean HR, high-frequency and low-frequency HRV, systolic and diastolic blood pressure, white matter hyperintensities and body-mass index to extract a set of latent variables reflecting variability in cardiovascular health across all individuals. The analysis used matlab *factoran.m* with default settings. Input variable distributions which deviated from Gaussian normality (1-sample Kolmogorov-Smirnov Test, p-value<0.05) were log-transformed (1-sample Kolmogorov-Smirnov Test, p-value > 0.05) (Fink, 2009).

### 2.7. Analytical approach

We performed both voxel-wise and component-based analyses using multiple linear regression (MLR) with robust fitting algorithm (matlab function fitlm.m). Voxel-level analysis was based on voxel-wise estimates across all imaging maps (RSFA, GM and ASL), while component-based analysis was based on component-wise estimates across all imaging components. We adopted a two-stage procedure for each RSFA voxel/component (Figure 1). In the first stage we used MLR with RSFA values for all individuals as dependent variable. The second stage correlated the residuals from each model with age.

In the first level models, independent variables included either cardiovascular health, CBF or grey matter measures and RSFA values as dependent variable. Covariates of no interest included gender, head motion and handedness. In the model with grey matter (model V, see below), the signal defined in the CSF mask was considered as a covariate of no interest to minimize the influence of non-morphological confounds in T1-weighted data (Bhogal et al., 2017; Ge et al., 2017; Tardif et al., 2017). Additional inclusion of total intracranial volume (TIV) did not change the principal results. Non-normally distributed variables were logarithmically or exponentially transformed to conform normality (Fink, 2009).

We constructed five models:

- Model 1: Covariates [of no interest]

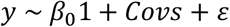
- Model 2: Covariates and cerebrovascular measures

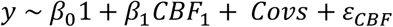
- Model 3: Covariates and cardiovascular measures

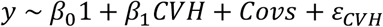
- Model 4: Covariates, cardiovascular and cerebrovascular measures

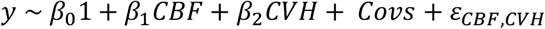
- Model 5: Covariates and grey matter volume measures

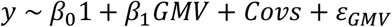

Note that the independent variables in Models 2, 4 and 5 included measures with voxel-specific information, i.e. RSFA values across subjects in a given voxel were predicted by the CBF/GM values for the corresponding voxel.

The residuals, ε, from each model were then used in a second-stage linear regression (i.e. correlational analysis) to estimate their association with age. Voxels where the residuals correlate with age (p<.05, FDR-corrected) indicate that the independent variables in first-stage model could not explain sufficiently the age-dependent variability in RSFA. Conversely, residuals not associated with age would suggest that the independent variables considered in the model are sufficient to explain age-dependent variability in RSFA.

This two-stage procedure was performed for each voxel of RSFA maps resulting in a statistics map for each model indicating the association between residuals and age. Statistical maps were corrected for multiple comparisons at p <0.05 (FDR-corrected). To further address multiple comparisons and voxel-voxel mapping between modalities, we performed complementary analysis where voxel-wise estimates of brain measures were substituted with subject-wise IC loadings, see Section 2.6.

We also tested whether the distribution of age-RSFA residuals correlations across all voxels formed differed from the predicted distribution under pure randomness. We constructed 5000 distributions of age-RSFA residual correlations across all brain voxels (D_Voxels_), where RSFA residuals were based either on a model with obseved RSFA values (D_Voxels_1) or permuted RSFA values (D_Voxels_2-5000). Distribution medians and distribution shapes were compared using Wilcoxon rank sum test and Kolmogorov-Smirnov test respectively. We performed a pair-wise comparison across all 5000 distribution shapes using Kolmogorov-Smirnov test, resulting in a distribution of 4999 similarity scores (D_Similarity_) between each D_Voxels_ with the remaining 4999 D_Voxels_. Next, we estimated the number of times (Np) the distribution of similarity for observed RSFA values (D_Similarity_1) is statistically different than the permuted distributions of similarities (D_Similarity_ 2-5000) using Wilcoxon rank sum test. The ratio Np/5000 provided a level of significance, e.g. a value < 0.05 suggested that the distribution of age-RSFA residual values is not as predicted by a model with pure randomness (at significance level p<0.05) and suggests an association between age and RSFA residuals. The procedure was applied separately for each of the five models across all brain voxels, as well as for different tissue types (cerebrospinal fluid, grey matter and white matter voxels with values above 0.4 in SPM’s tissue probability maps).

## 3. Results

### 3.1. Main and age effects of RSFA, CBF and CVH

#### 3.1.1. Resting state fluctuation amplitudes (RSFA)

Whole group voxel-wise analysis revealed relatively high RSFA values (relative to the average across the brain) across all individuals in the frontal orbital, inferior frontal gyrus (IFG), dorsolateral prefrontal cortex (dlPFC), superior frontal cortex, anterior and posterior cingulate, and lateral parietal cortex (Figure 2a). With respect to aging, we observed age-related decreases in RSFA in the bilateral IFG, bilateral dlPFC, bilateral superior frontal gyrus, primary visual cortex, cuneus, precuneus, posterior and anterior cingulate, superior temporal gyrus, medial parietal cortex, and lateral parietal cortex (Figure 2b). Regions in the proximity of frontal white matter, cerebrospinal fluid and large vascular vessels showed a significant increase of RSFA values as a function of age.

**Figure 2.**
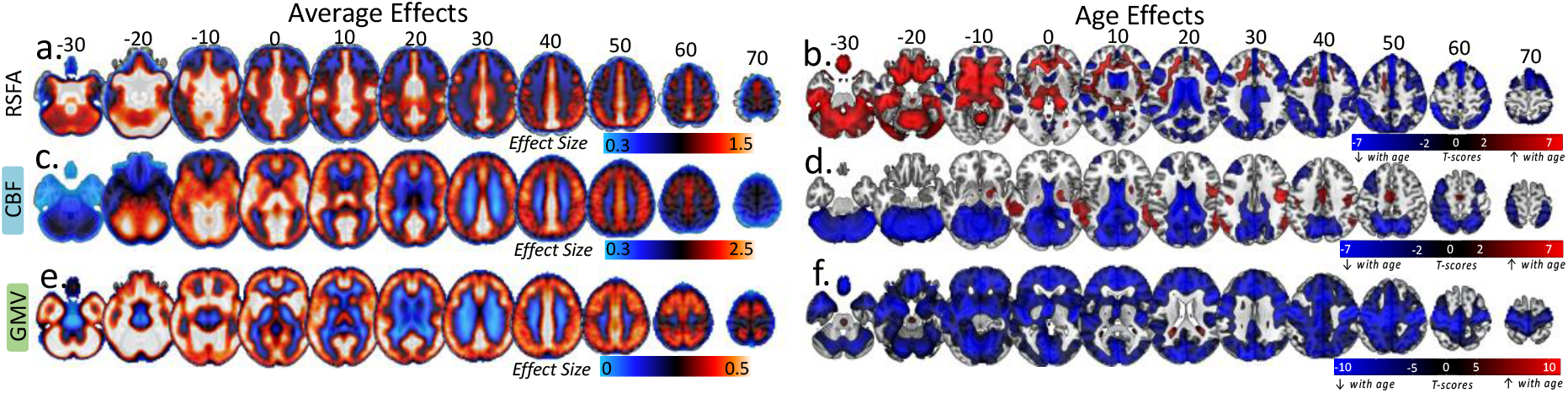
Average RSFA, CBF and Grey Matter Volume and the effects of age on each modality (SPM{beta} and SPM{t} maps respectively)

#### 3.1.2. Cerebral blood flow (CBF)

Whole group voxel-wise analysis revealed a pattern of relatively high cerebral blood flow across all individuals in cortical and subcortical brain areas with high perfusion and metabolism properties (Figure 2c) including caudal middle-frontal, posterior cingulate, pericalcarine, superior temporal and thalamic regions. Moderate to low CBF values in the superior-parietal and inferior-frontal areas of the cortex (Figure 2c, every 10 axial slices from −30 to 70) may reflect the axial positioning of the partial brain coverage sequence used in the study. With respect to aging, we observed age-related reductions in CBF in the bilateral dorsolateral prefrontal cortex, lateral parietal cortex, anterior and posterior cingulate, pericalcarine and cerebellum (Figure 2c). In addition, we observed age-related CBF increase in regions susceptible to individual and group differences in in arterial transit time biasing the accuracy of CBF estimation, including middle temporal gyrus (Mutsaerts et al., 2017).

#### 3.1.3. Grey matter volume (GMV)

We identified significant whole-group effects across all grey matter voxels (Figure 2e). In addition, there was a widespread age-related decrease in GMV, in bilateral temporal lobes, bilateral prefrontal, middle and superior frontal areas, bilateral medial occipital areas, cerebellum, and subcortical areas including thalamus, caudate and putamen (Figure 2f), consistent with previous reports (Mohajer et al., 2020; Peelle et al., 2012; Tsvetanov et al., 2019).

#### 3.1.4. Cardiovascular health (CVH)

An exploratory factor analysis with principal component analysis indicated a three-factor structure of the cardiovascular health and risk measures. Factor 1 loadings indicated a factor expressing variability in blood pressure measures, where individuals with higher subject scores had larger systolic and diastolic pressure (Figure 3). Subjects scores did not correlate with age (r = +.061, p=.328), indicating that variability in blood pressure was not associated uniquely with aging over and above their contribution to other factors in the analysis. Factor 2 was mainly expressed by heart rate and HRV measures, where individuals with high subject scores had low resting pulse and high HRV metrics. Subject scores were correlated negatively with increasing age (r = −.417, p<.001), consistent with findings of age-related decrease in HRV (Figure 3). Finally, Factor 3 was expressed negatively by HRV and positively by WMH and systolic blood pressure, indicating that individuals with high subjects scores were more likely to have high burden of WMH, high systolic blood pressure and low HRV (Figure 3). Subject scores were associated positively with age (r = +.713, p<.001), suggesting that a portion of the age-related decrease in HRV is coupled with increase in WMH and systolic blood pressure.

**Figure 3.**
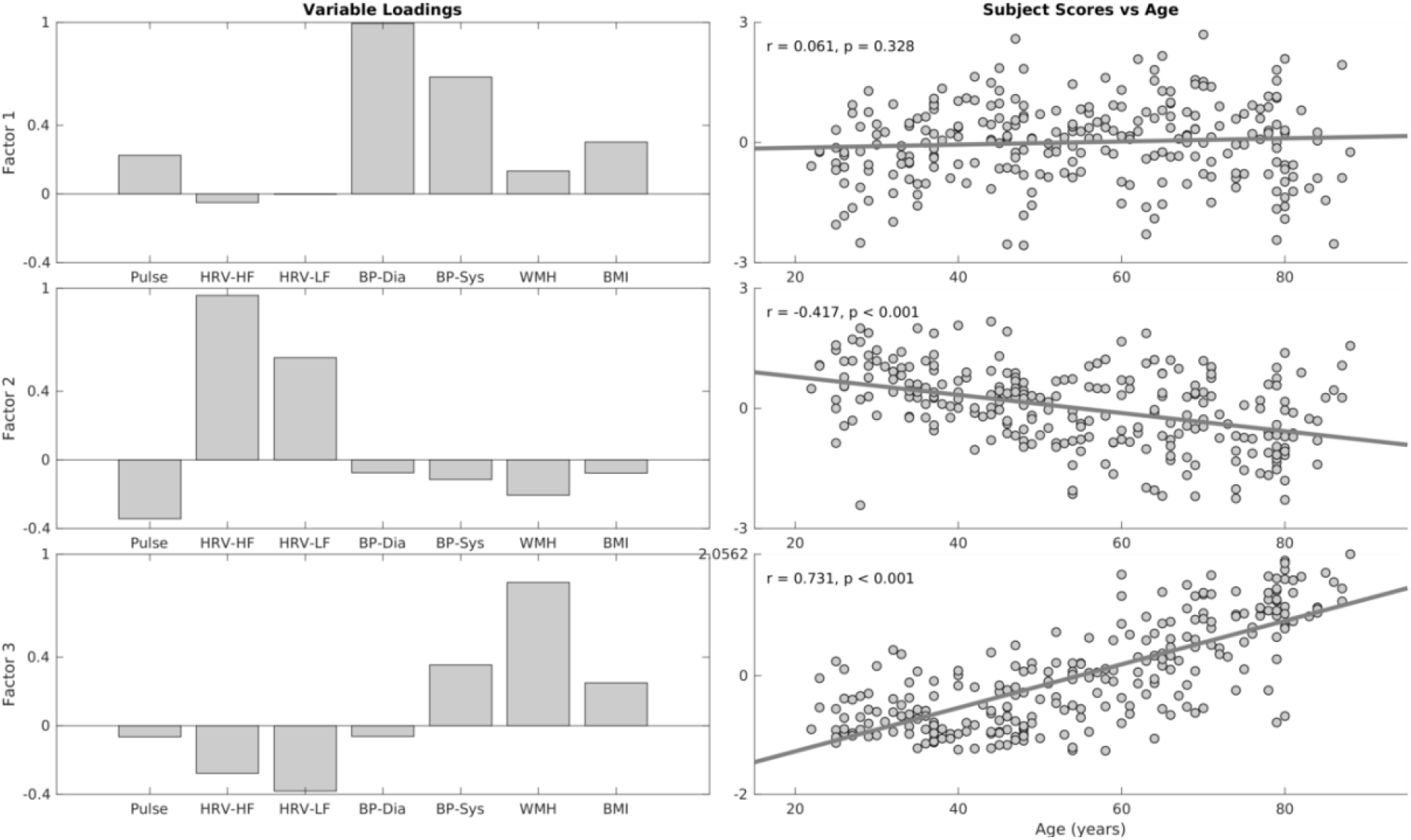
Variable loadings (left column) and association between age and subjects scores for three factors resulting from factor analysis on cardiovascular risk variables. Pulse – mean heart rate, HRV-HF – high-frequency heart rate variability, HRV-LF – low-frequency high rate variability, BP-Dia – diastolic blood pressure, BP-Sys – systolic blood pressure, WMH – white matter hyperintensities, BMI – body-mass index

### 3.2. Correlations between Age and RSFA residuals

#### 3.2.1. Voxel-based analysis

##### Covariates of no interest only (Model I)

The whole-group voxel-wise analysis of RSFA maps revealed brain regions with high vascular reactivity including frontal orbital, inferior frontal gyrus, inferior frontal gyrus, dorsolateral prefrontal cortex, superior frontal cortex, anterior and posterior cingulate, and lateral parietal cortex. We observed age-related decrease in RSFA in the bilateral inferior frontal gyrus, bilateral dorsolateral prefrontal cortex, bilateral superior frontal gyrus, primary visual cortex, cuneus, precuneus, posterior and anterior cingulate, superior temporal gyrus, medial parietal cortex, and lateral parietal cortex. In addition, we observed age-related decrease in RSFA in the proximity of ventricles and large vascular vessels.

##### Controlling for Cerebrovascular Factors (Model II)

We observed significant correlations between age and the RSFA residuals after controlling for subject variability in CBF and covariates of no interest at an FDR-adjusted p-value of 0.05 (Figure 4, model II). The spatial extent and the size of the statistical maps were similar to the analysis with RSFA residuals after controlling for covariates only (Figure 2d and Figure 4, model I), suggesting that CBF does not fully explain variability in RSFA.

**Figure 4.**
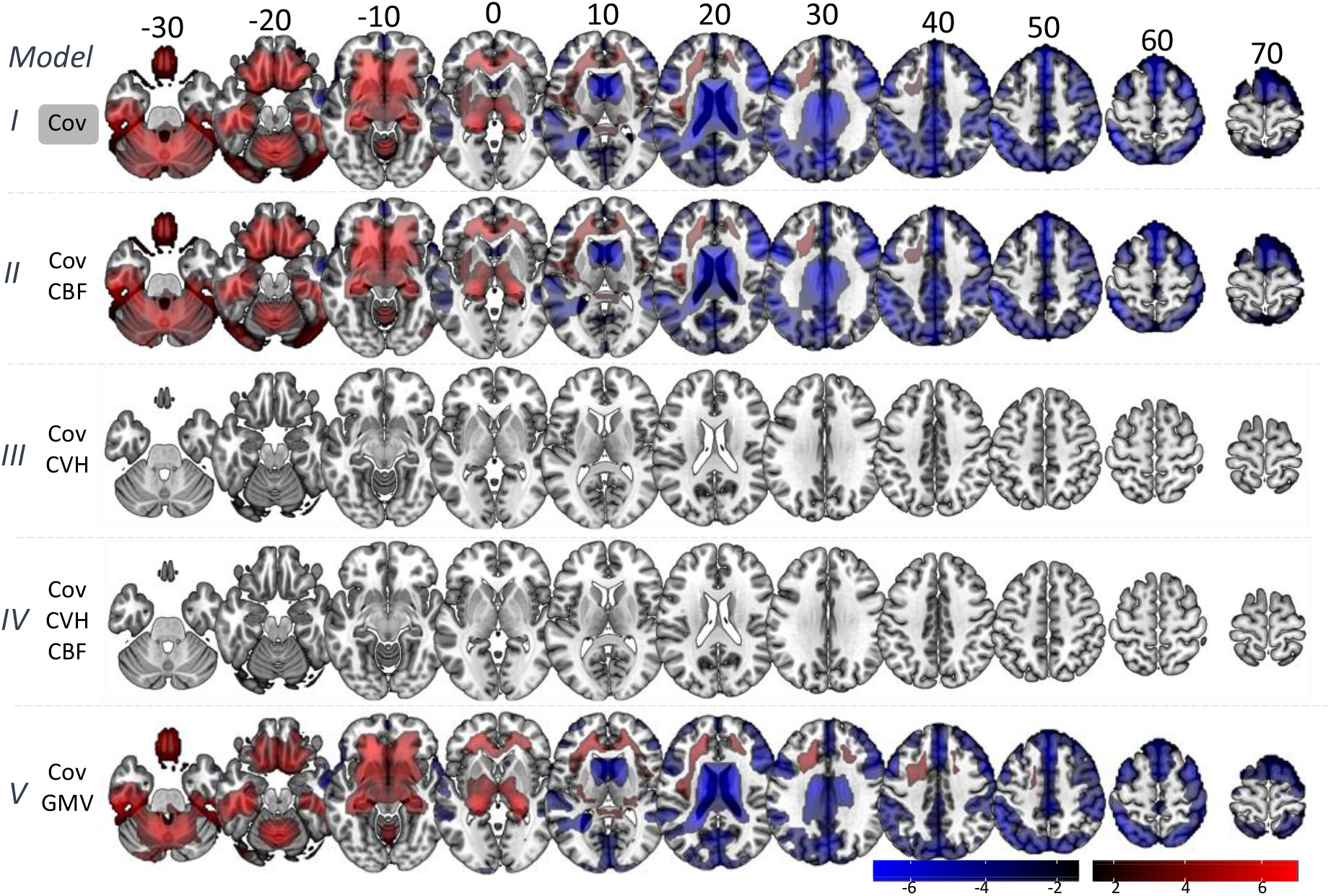
Voxel-wise associations between age and RSFA residuals after controlling for: covariates only (Cov, Model I); Cov and cerebral blood flow (CBF, Model II); Cov and cardiovascular health (CVH, Model III); Cov, CBF and CVH (Model IV); and Cov and grey matter volume (GMV).

##### Controlling for Cardiovascular Factors (Model III)

We observed no significant correlations between age and the RSFA residuals after controlling for variability in CVH and covariates of no interest at an FDR-adjusted p-value of 0.05 (Figure 4, model III), suggesting that CVH can explain sufficiently age-dependent variability in RSFA, at least at the level of statistically-corrected voxels.

##### Controlling for Cardiovascular and Cerebrovascular Factors (Model IV)

We observed no significant correlations between age and the RSFA residuals after controlling for variability in CVH, CBF and covariates of no interest at an FDR-adjusted p-value of 0.05 (Figure 4, model IV), suggesting that CVH and CBF together explain sufficiently age-dependent variability in RSFA.

##### Controlling for Grey Matter Volume (Model V)

We observed significant correlations between age and the RSFA residuals after controlling for grey matter volume (GMV) and covariates of no interest at an FDR-adjusted p-value of 0.05 (Figure 4, model V), suggesting that GMV does not adequately explain variability in RSFA, at the voxel-wise level.

#### 3.2.2. Distribution-based analysis

The medians of observed and permuted data did not differ significantly (p>.1 for all five models). In terms of the distributions, the level of statistical significance decreased after controlling for cardiovascular, cerebrovascular and GMV signals (p <.001, p<.001, p=.015, and p<.001 for models 1, 2, 3 and 5 respectively), see Table 2. The model considering jointly cardiovascular and cerebrovascular signals (model 4) indicated a difference in the distribution of observed and permuted data (p = 0.016), reflecting a small level of correlation between age and RSFA residuals in some voxels. It is unclear whether the signal originated in a particular tissue type, so we repeated the permutation approach for each tissue type separately (Table 2). For models 1, 2 and 5 the RSFA residuals were associated with age across all three tissue types, suggesting that variability in cerebrovascular and grey matter cannot account fully for the effects of age on RSFA in all tissue types. However, the models controlling for cardiovascular health (Models 3 and 4) were not significant for grey matter and white matter tissue. The analysis on CSF voxels was highly significant suggesting that any potential age-related effects on RSFA not captured by cardiovascular and cerebrovascular signals on voxel-level are focal to CSF areas, rather than grey matter or white matter.

**Table 2.** Evaluation of the difference in distribution shape across voxels in the whole brain, as well as voxels within grey matter (GM), white matter (WM) and cerebrospinal fluid (CSF) areas. Tests showing no difference in the distributions at uncorrected p-value 0.05 are indicated by n.s.

| Model | Whole Brain | GM | WM | CSF |
| --- | --- | --- | --- | --- |
| 1 | <0.001 | <0.001 | 0.009 | <0.001 |
| 2 | <0.001 | <0.001 | 0.009 | <0.001 |
| 3 | 0.015 | n.s. | 0.048 | 0.005 |
| 4 | 0.016 | n.s. | 0.039 | 0.007 |
| 5 | <0.001 | <0.001 | 0.004 | <0.001 |

#### 3.2.3. Component-based analysis

CVH signals sufficiently explained variance in RSFA, in the voxel-based analysis (after FDR correction for multiple comparisons) and in grey-matter areas in distribution-based analysis. This was not the case for CBF or GMV in the voxel-based analysis, as well as for CVH in CSF regions in distribution-based analysis. However, this might reflect limitations of these analyses to separate spatially overlapping sources of signal with different aetiology and the large number of comparisons (see Methods). Therefore, we used independent component analysis to decompose each imaging modality to a small number of spatially-independent components and test their ability to explain variance of RSFA.

Figure 5 shows the decomposition of the RSFA, CBF and GMV datasets with 18, 13 and 16 number of components, respectively, according to the MDL criterion (Li et al., 2007). The spatial maps of the components and the between subject-correlations of loading values revealed patterns of signal from grey matter, white matter, cerebrospinal fluid and vascular aetiology (Figure 5), which were highly consistent with voxel-wise analysis (Figure 2), previous reports of RSFA (Tsvetanov et al., 2015) and structural data (Eckert et al., 2010; K. Liu et al., 2017).

**Figure 5.**
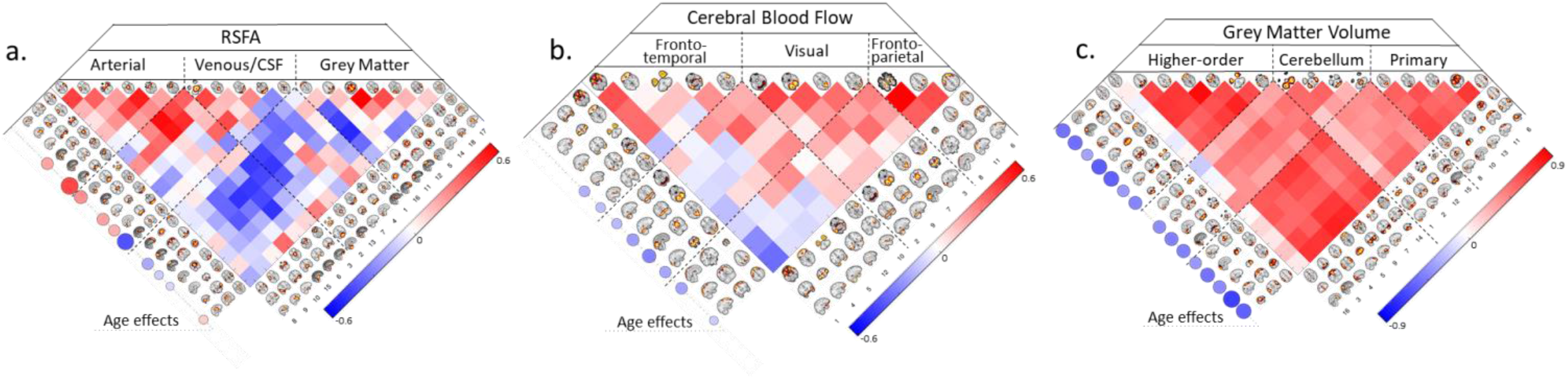
Inependent component analysis spatial maps and correlation between subject loadings for RSFA (a), cerebral blood flow (b) and grey matter volume (c) datasets. The relationships between age and IC loadings are shown circles on the left hand-side of each correlation matrix, FDR-adjusted p-value of 0.05.

The effects of ageing on the independent components loadings was consistent with the voxel-level analysis. Specifically, RSFA components with vascular ethology indicated an age-related increase in the loading values, while ICs confined within grey matter areas showed age-related decrease in the loading values (Figure 5a, left side of the panel). Several CBF components demonstrated age-related decrease in loading values, including inferior frontal gyrus, superior frontal gyrus, cuneus, precuneus, lateral occipital cortex and motor cortex (Figure 5b, left side of the panel). All but one GMV component in the cerebellum demonstrated age-related decrease in loading values consistent with brain-wide atrophy in ageing (Figure 5).

Next, we turn to the correlations between age and residuals of the RSFA ICs. We focused on ICs that showed age-related differences in the subject loading values (10 out of 18), after controlling for CBF IC loading values, GMV IC loading values or CVH factor loadings (Figure 6).

**Figure 6.**
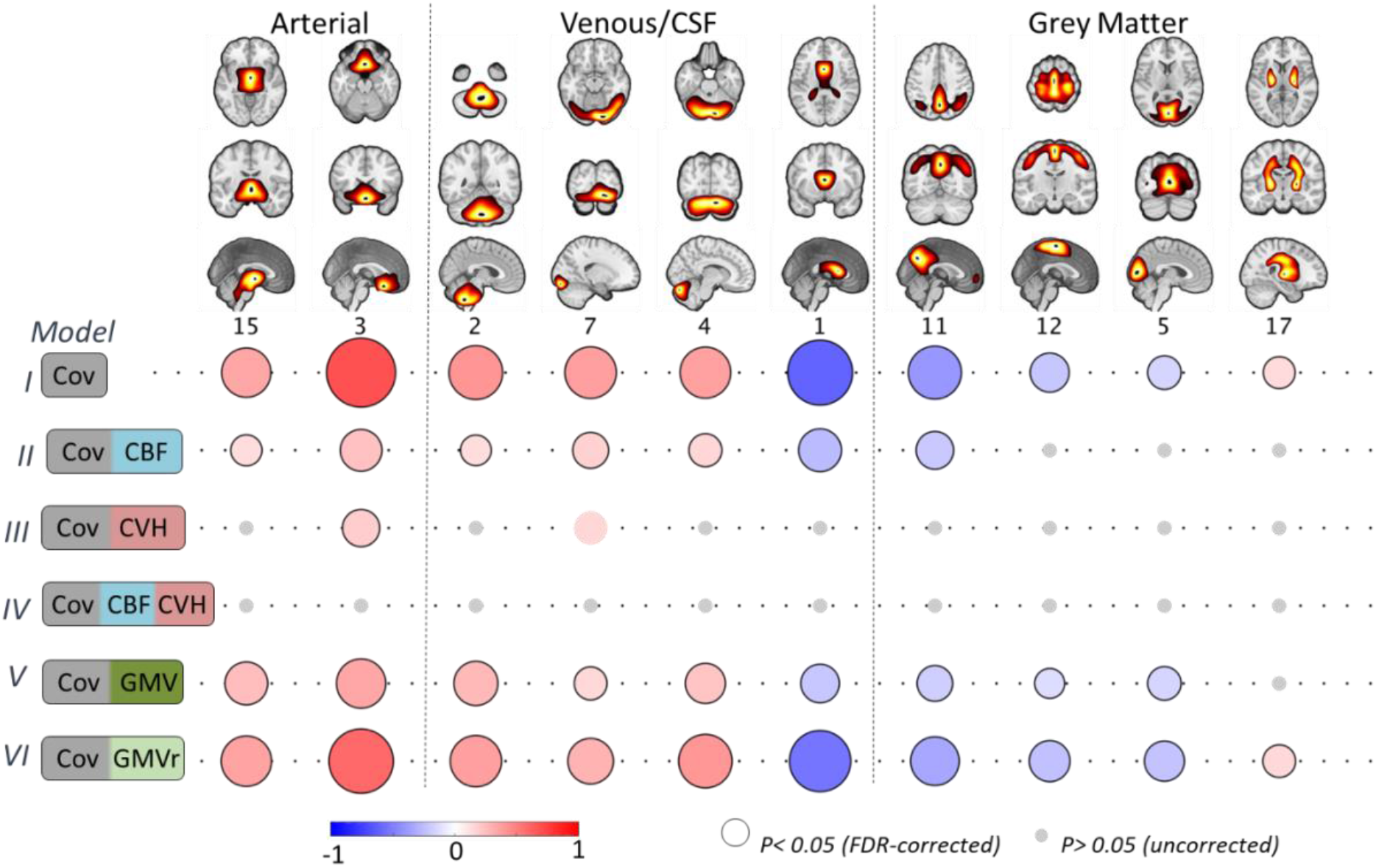
Component-based associations between age and RSFA residuals after controlling for: covariates only (Cov, Model I); Cov and cerebral blood flow (CBF, Model II); Cov and cardiovascular health (CVH, Model III); Cov, CBF and CVH (Model IV); Cov and grey matter volume (GMV); and Cov and grey matter volume residuals (GMVr) after controlling for the effects of CVH (see text). Grey circles denote uncorrected p-value >0.05, circles without black outline denote uncorrected p<0.05 and circles with black outline denote FDR-adjusted p-value at 0.05.

##### Controlling for Cerebrovascular Factors (Model II)

The associations between age and RSFA residuals after controlling for CBF loading values were weaker in vascular ICs and abolished in GM ICs compared to the analysis with covariates only (Figure 6, Model I vs Model 2). Unlike in the voxel-based analysis, this ICA approach suggests that CBF does explain some age-related variability in RSFA across many networks, especially those in GM areas, which may be due to reduced number of comparisons and improved characterisation of sources of signals in RSFA and CBF data using ICA.

##### Controlling for Cardiovascular Factors (Model III)

After controlling for differences in CVH, RSFA residuals in two ICs (IC3 and IC7) were correlated with age (uncorrected p-value at 0.05 significance level), although to a lesser extent compared to the analysis with covariates only (Model III vs Model I), indicating that CVH can explain age-dependent variability in most, but not all, RSFA ICs.

##### Controlling for Cerebrovascular and Cardiovascular Factors (Model IV)

We observed no significant correlations between age and the RSFA residuals after controlling for variability in CVH and CBF (even at an uncorrected p-value of 0.05, see Figure 6), suggesting that together, CVH and CBF can explain age-dependent variability in RSFA.

##### Controlling for Grey Matter Volume (Model V)

RSFA ICs adjusted for GMV ICs demonstrated reduced correlations between RSFA and age (particularly RSFA ICs of grey matter territories), indicating that age-related differences in RSFA ICs can be partly explained by grey matter atrophy.

##### Controlling for Grey Matter Volume independent of Cardiovascular Factors

Some degree of association between age differences in RSFA and grey matter atrophy is expected given cardiovascular health has been linked to brain-wide atrophy (Gu et al., 2019; Srinivasa et al., 2016) and T1-weighted data is confounded by non-morphological signals (Bhogal et al., 2017; Ge et al., 2017; Tardif et al., 2017). Therefore, to test whether the effects of brain atrophy on RSFA were independent of the effects of CVH on brain atrophy, we controlled for the effects of CVH in GMV ICs. Then we used the GMV residuals after fitting CVH to GMV IC loadings (i.e. GMV orthogonalised with respect to CVH) to estimate RSFA residuals and subsequently their correlation with age (Figure 6, Model 6). The effects between RSFA residuals and age in Model 6 were similar to Model 1, suggesting that GMV differences independent of CVH were not correlated to differences in RSFA.

## 4. Discussion

The principle result of this study is to confirm the suitability of resting state fluctuation amplitude (RSFA) to quantify vascular influences in BOLD-based fMRI signals, and to demonstrate that the age effects on RSFA reflect variability in vascular factors rather than neuronal factors. We demonstrate that the effects of age on RSFA can be sufficiently captured by the joint consideration of cardiovascular (based on ECG, BP, WMH and BMI measures) and cerebrovascular factors (CBF from ASL). Variance in brain atrophy (GM volume Figure 6) and neuronal activity (Kumral et al., 2019; Tsvetanov et al., 2015) do not explain unique relationship between RSFA and age. This means that RSFA is a suitable measure for differentiating between vascular and neuronal influences on task-based BOLD signal. Without modelling the age-related differences in cardiovascular and cerebrovascular factors, changes in ‘activity’ based on BOLD-fMRI could be misinterpreted, thereby undermining conceptual advances in cognitive ageing.

### Cardiovascular factors and age-differences in RSFA

We used factor analysis to estimate cardiovascular health from a wide range of cardiovascular measures (Varadhan et al., 2009; Wardlaw et al., 2014). Our three factor solution resembled previous reports (Chen et al., 2000; Goodman et al., 2005; Khader et al., 2011; Mayer-Davis et al., 2009), with two factors associated with blood pressure and heart rate variability (factors 1 and 2, respectively). A third factor expressed white matter hyperintensities, blood pressure, heart rate variability and body-mass index, suggesting a cerebrovascular origin.

These three factor indices of cardiovascular health explained most of the age-related variability in RSFA, leaving little to no associations between age and RSFA residuals in grey matter regions (after controlling for these cardiovascular signals). This suggests that differences in cardiovascular health mediate most of the age effects on RSFA (Tsvetanov et al., 2015). Interestingly, each CVH factor was associated with a distinct spatial RSFA pattern (Supplementary Figure 2) and collectively provided additional explanatory value for the overall age-differences in brain-wide RSFA. Next, we turn to neural and cerebrovascular contributions to BOLD.

### Cerebrovascular signals and age-differences in RSFA

Our measure of cerebrovascular function was based on cerebral blood flow estimates from a common perfusion-based ASL sequence. Here, we refer to cerebrovascular function as an umbrella term of physiological alterations in the neurovascular unit including resting CBF, cerebrovascular reactivity, cerebral autoregulation and pulsatility. The observed average, gender and age effects were consistent with previous reports. The age effects on CBF values were in agreement with previous reports (Chen et al., 2011; Zhang et al., 2018), with decreases mainly found in regions that are associated with high perfusion and metabolic demand, including precuneus, cuneus, prefrontal cortices and cerebellum. The mechanisms underlying the observed CBF decrease across the adult lifespan is a subject of continuous debate between structural and physiological alterations of the neurovascular unit (Girouard and Iadecola, 2006; Tarumi and Zhang, 2018; Tsvetanov et al., 2020). We also observed age-related increase in CBF in temporal regions, which may reflect macro-vascular artifacts that are common to arterial spin labelling findings (Detre et al., 2012; Mutsaerts et al., 2017) due to prolonged arterial transit time with ageing (Dai et al., 2017). This nonspecific nature of resting CBF signal changes during ageing is particularly problematic for fMRI BOLD studies, since differences in physiology on that level may confound the interpretation of the BOLD signal as a surrogate measure of evoked neural activity (Whittaker et al., 2016).

Compared to voxel-wise estimates, our component-wise CBF values captured better the age-related effects of RSFA, especially in grey matter areas (see below on differences between voxel-wise and component-wise analysis). Nevertheless, neither the voxel-wise nor component-based analysis of CBF values could explain sufficiently the effects of age on RSFA, suggesting that RSFA may not be attributed exclusively to sources of signal linked to cerebrovascular function (Garrett et al., 2017; Liu et al., 2012). There was a positive correlation between resting CBF and RSFA in brain areas typically associated with high blood perfusion and metabolic demands, including cuneus, precuneus, intraparietal sulcus, inferior temporal cortices, dorsolateral prefrontal cortex and anterior cingulate (Supplementary Figure 3). But, we also observed negative associations between RSFA and CBF in inferior brain areas, mainly close to vascular territories i.e. the higher the RSFA the lower the CBF values were in these regions (Supplementary Figure 3). This may reflect the dominance of pulsatility influences in RSFA signals near vascular territories and the CSF (for more information see section: Spatial distribution and age effects on RSFA). This may have adverse effects on tissue perfusion in neighboring areas (Tarumi et al., 2014). The coexistence between positive and negative relationships between RSFA and CBF measures in our study explains previous observations of a varied direction in the relationship between these measures across regions for groups and individuals with differences in vascular health (Garrett et al., 2017).

### Joint effect of cardiovascular and cerebrovascular factors

The joint consideration of cardiovascular and CBF measures fully explained the (significant) effects of age on RSFA in grey matter regions, despite their differential association with ageing (Zlokovic, 2011). This suggests that RSFA can normalize BOLD fMRI for both cardiovascular and cerebrovascular factors as highly reliable and temporally stable measurement compared to current standard approaches to normalize BOLD fMRI (eg. hypercapnia) (Golestani et al., 2015; Lipp et al., 2015). Lower reproducibility in “gold standard” approaches could be due to susceptibility of cerebrovascular measures to short-term variable physiological modifiers (e.g. caffeine, nicotine, time of the day, drowsiness) (Clement et al., 2018). The high reproducibility of RSFA in healthy adults could come from the additional contribution of short-term but stable cardiovascular health signals (e.g. heart condition or white matter hyperintensities), which are independent of cerebrovascular factors. RSFA reflects both cardiovascular and cerebrovascular signals, which are associated with distinct spatial patterns (see section Spatial distribution and age effects on RSFA). RSFA can help dissociate age-related differences in cardiovascular, cerebrovascular and neural function in task-based BOLD signal, which is important for using fMRI to understand the mechanisms of cognitive aging.

### Grey matter volume and age-differences on RSFA

Voxel-wise and component-based analyses indicated weak associations between age-differences in RSFA and grey matter volume, which were abolished after adjusting for variability in cardiovascular health. Interestingly, the strongest effects were at the boundaries between grey matter and other tissue types (white matter and CSF), rather than deep cortical areas (Supplementary Figure 3). The spatial pattern of the effects for cortical areas was similar to those observed between CBF and RSFA measures. There was a positive relationship between RSFA and grey matter volume in the precuneus, intraparietal sulcus, dorsolateral prefrontal cortex and dorsal anterior cingulate; which could reflect the cerebrovascular component of the RSFA signal (see above). In addition, the cerebellum and subcortical areas near vascular territories showed negative associations, i.e. individuals with less grey matter volume had larger RSFA values, likely reflecting the cardiovascular components of the RSFA signal (see below, Spatial distribution and age effects of RSFA). Importantly, there were no associations between RSFA and grey matter volume after adjusting for cardiovascular health. This is suggestive of an indirect association between RSFA and grey matter volume introduced by cardiovascular effects on brain-wide atrophy (Gu et al., 2019; Srinivasa et al., 2016) and other non-morphological confounds in T1-weighted data (Bhogal et al., 2017; Ge et al., 2017; Tardif et al., 2017). The lack of evidence for an association between age-related effects on RSFA and brain atrophy after adjusting for cardiovascular health is consistent with previous reports using direct physiological measures of neural activity (MEG and EEG): no age-related associations between RSFA and neuronal indices were detected (Kumral et al., 2019; Tsvetanov et al., 2015). Furthermore, potential age-related associations between RSFA and cognitive function are fully explained by cerebrovascular risk factors, such as WMH burden (Millar et al., 2020). Taken together these findings suggest that the age-related differences in BOLD signal variability at resting state are unlikely to be of neuronal origin beyond the effects of age on various types of vascular signals.

### Spatial distribution and age effects on RSFA

The voxel-wise and component-based analysis of RSFA maps reveal brain regions with high vascular reactivity (Di et al., 2012; Kalcher et al., 2013; Kannurpatti et al., 2011; Liu et al., 2013; Mueller et al., 2013; Yezhuvath et al., 2009), and accord with previous studies of average and age-effects on RSFA (Golestani et al., 2016; Lipp et al., 2015; P. Liu et al., 2017; Tsvetanov et al., 2015). These patterns of spatially distinct cortical areas might reflect segregation of cortical tissue composition, e.g. delineation on the basis of vascular density and metabolic demands in areas with cyto- or myeloarchitectonic differences (Annese et al., 2004; Fukunaga et al., 2010; Geyer et al., 2011; Glasser and Van Essen, 2011). The age-related increase in RSFA in areas with vascular, WM and CSF partitions may reflect the impact of vascular pulsatility downstream of cerebral arteries due to wall stiffening of blood vessels (Robertson et al., 2010; Webb et al., 2012), which may influence BOLD signal variability in neighboring brain tissue (Lee and Oh, 2010; O’Rourke and Hashimoto, 2007; Tarumi et al., 2014; Viessmann et al., 2017). The pulsatility can influence signal in white matter and cerebrospinal fluid areas (Makedonov et al., 2013; Tarumi et al., 2014; Theyers et al., 2018; Viessmann et al., 2019). In addition, it is also possible that the RSFA signal in one area of the brain captures the presence of multiple sources of signal with different aetiology. For example, the observed signal in one CSF voxel may be a mixture of signals coming from fluctuations in resting CBF in neighboring vascular territories and pulsatility influences in the perivascular space. Spatially overlapping sources of signal might be difficult to detect and dissociate using a univariate approach. This motivates the use of multivariate data-driven approaches, as highlighted by our findings. In sum, this suggests that RSFA reflects different types of vascular signals with distinct spatial patterns in terms of signals with cerebrovascular origin in grey matter regions, and those with cerebro- and cardio-vascular origin in other parts of the brain.

### Limitations and future directions

There are limitations to the current study. In terms of cardiovascular health, there may be more important measures that were not present in the CamCAN sample. Moreover, the analysis of heart rate variability estimates was based on normal-to-normal beats (Vest et al., 2018). The difference between NN- and RR-beat analysis is that the former considers the detection and exclusion of segments and participants with atrial fibrillation and other abnormal beats. While NN-beat analysis optimises the detection of unbiased estimates of cardiovascular health, it also precludes sensitivity to potential effects of arrhythmia and abnormal heart beats on RSFA in our analysis, which might be relevant to regions susceptible to pulsatility effects (Webb and Rothwell, 2014).

In terms of cerebrovascular signals, the use of ASL-based CBF measurements could be complemented with individual-based arterial transit time measurement in order to improve the accuracy of ASL imaging in older populations (Dai et al., 2017). There are also other means to assess cerebrovascular function, including cerebrovascular reactivity, including CO2-inhalation-induced hypercapnia (Liu et al., 2019), breath-hold-induced hypercapnia (Handwerker et al., 2007; Mayhew et al., 2010; Riecker et al., 2003; Thomason et al., 2007, 2005), hyperventilation-induced hypocapnia (Bright et al., 2009; Krainik et al., 2005), and venous oxygenation (Liau and Liu, 2009; Lu et al., 2010; Restom et al., 2007) and it is possible that these might reveal effects in RSFA where ASL-based CBF does not. Future studies should explore the utility of additional estimates from resting ASL-based CBF data to complement CBF quantification. For instance, little is known about whether resting CBF variability, which is statistically similar to RSFA, is sensitive to cerebrovascular reactivity and other vascular origins (Robertson et al., 2017). The ease, safety and tolerability of RSFA across the lifespan yields a considerable advantage for population and clinical studies.

Similar to the CBF analyses, the GMV findings generalized across voxel-wise and component-based analysis, but the component-based analysis seemed to be more sensitive to the age effects on RSFA in both CBF and GMV datasets. The greater generalization across datasets with independent component analysis than voxel-based analysis may reflect several factors (Calhoun and Adali, 2008; Passamonti et al., 2019; Sui et al., 2012), e.g. reducing the burden of multiple comparisons, pooling information across multiple voxels with similar profiles, separating sources of signal with different etiology but with overlapping topologies and possibly improving the spatial correspondence across imaging modalities with different spatial scales, sequence parameters and signal properties. Therefore, the use of component-based analysis in studies comparing approaches for normalization of physiological signals may improve understanding the nature of the signal and the extent to which these neuroimaging modalities are related to one another.

In the current study, RSFA was estimated from resting state fMRI BOLD data prior to collection of other task-based fMRI scanning as in previous validation studies of RSFA (Kannurpatti and Biswal, 2008; Tsvetanov et al., 2015). Other means of RSFA-like estimates have been proposed for scaling BOLD activation data using fMRI BOLD data at different non-resting cognitive states, e.g. during task periods (Kazan et al., 2016) or fixation-/resting-periods succeeding task periods (Garrett et al., 2017). Given that short periods of cognitive engagement can modulate the BOLD signal in a subsequent resting state scan (Sami et al., 2014; Sami and Miall, 2013), future studies are required to generalise our findings to RSFA-like estimates derived from other types of fMRI BOLD acquisition.

Finally, this study has focussed on the effects of aging, but other studies aiming to understand individual differences or drug effects in fMRI BOLD might be affected in a similar manner. Therefore, future studies should consider the origins of the signal contributing to RSFA (cerebrovascular vs cerebrovascular) and more broadly their influence in fMRI BOLD imaging studies. In the light of increasing evidence of the role of cardiovascular and cerebrovascular factors in maintaining cognitive function, future studies might even consider RSFA as a predictor, rather than just as a covariate of no interest, when modelling the effects of interest (e.g. age or performance). Furthermore, while the proposed approach is based on plausible neurophysiology that can be used to evaluate its contribution to cognitive function, future studies could improve absolute quantification of neural function together with its integration with deoxyhaemoglobin-dilution-based modelling (Davis et al., 1998; Hoge et al., 1999a, 1999b), haemodynamic response function modelling (West et al., 2019), generative modelling (Friston et al., 2003; Jafarian et al., 2020; Tsvetanov et al., 2016) and model-free decomposition (Bethlehem et al., 2020; Campbell et al., 2015; Samu et al., 2017; Tsvetanov et al., 2018) of fMRI BOLD data.

### Concluding remarks

Cardiovascular and cerebrovascular signals together predict the age differences in RSFA, establishing RSFA as an important marker that can be used to accurately separate vascular signals from neuronal signals in the context of BOLD fMRI. We propose that RSFA is suitable to normalize BOLD, and control for differences in cardiovascular signals. This is particularly relevant to the research in neurocognitive aging, and may reduce selection bias, for example by permitting the inclusion of individuals with a wider range of hypertension, cardiovascular conditions or comorbidity. The use of RSFA as a mechanism to adjust for confounds in BOLD-fMRI, or as a predictor, will allow the development of better models of ageing and age-related disorders (Cabeza et al., 2018; Tsvetanov et al., 2018).

## 5. Acknowledgements

This work is supported by the British Academy (PF160048), the Guarantors of Brain (101149), the Wellcome Trust (103838), the Medical Research Council (SUAG/051 G101400; and SUAG/046 G101400), European Union’s Horizon 2020 (732592) and the Cambridge NIHR Biomedical Research Centre.

## 9. Supplementary Figures

**Supplementary Figure 1.**
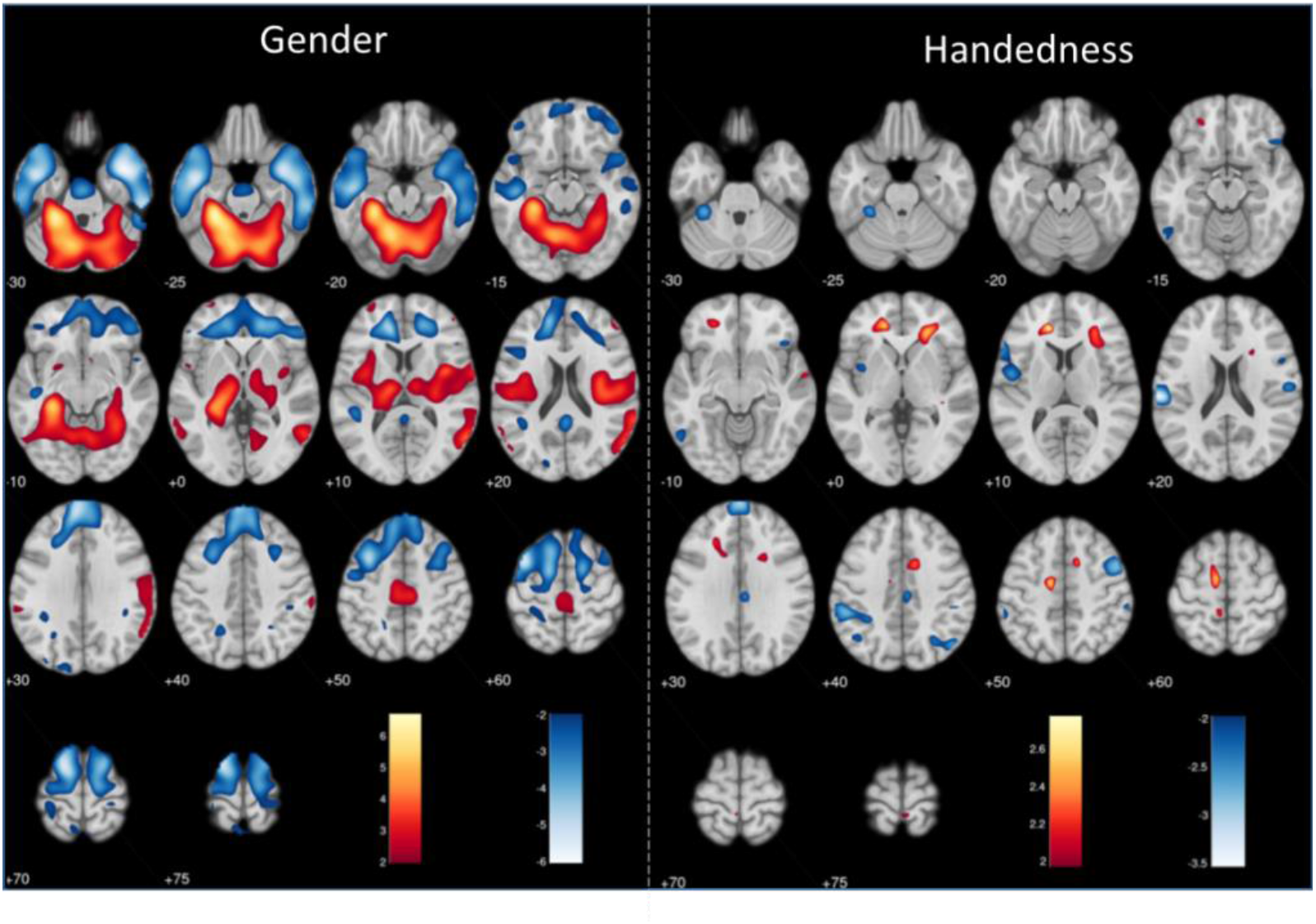
Voxel-wise associations between RSFA and covariates of no interests (gender – left panel and handedness – right panel), Model I. Maps are thresholded at uncorrected p-values of 0.05 for more complete description of the spatial represnation.

**Supplementary Figure 2.**
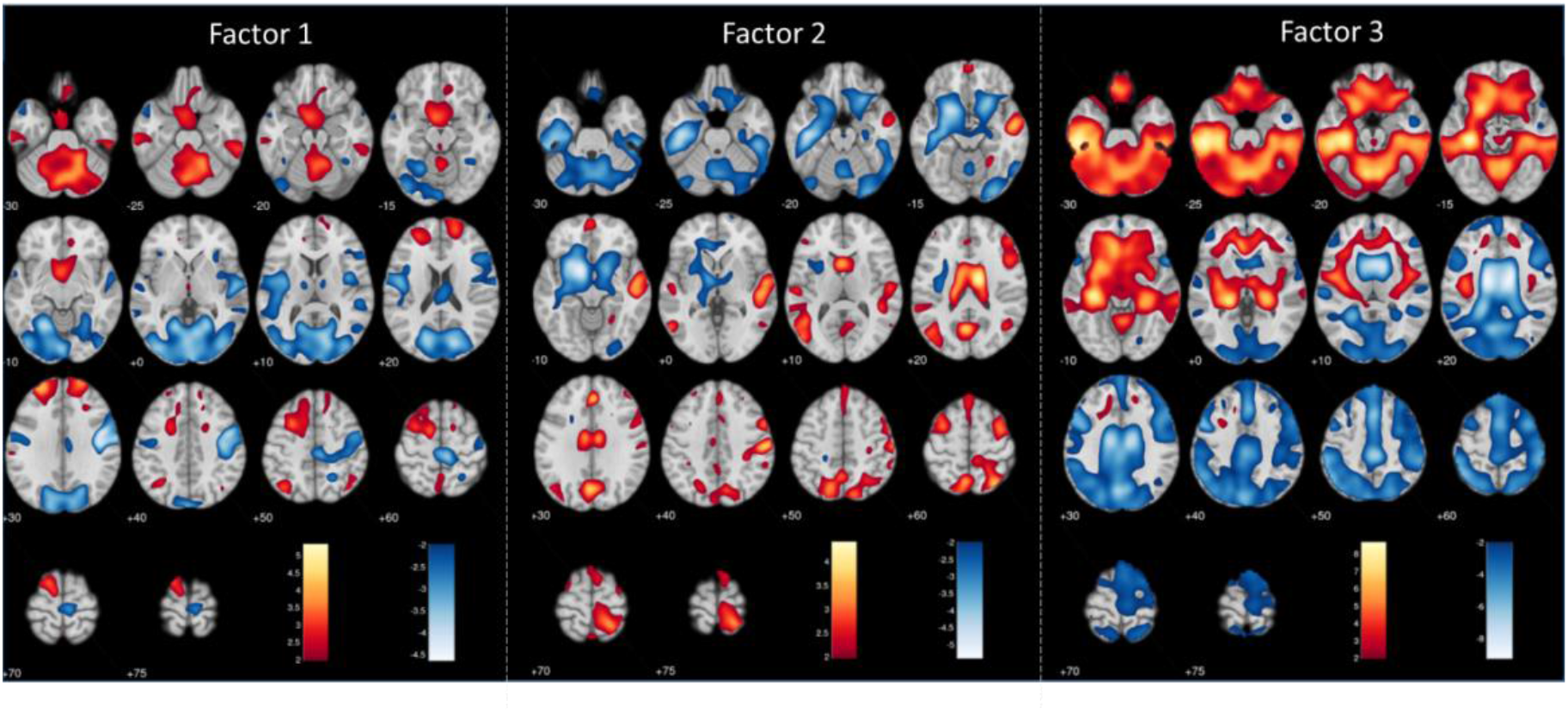
Voxel-wise associations between RSFA and three factors of cardiovascular health (Model III). Maps are thresholded at uncorrected p-values of 0.05 for more complete description of the spatial represnation.

**Supplementary Figure 3.**
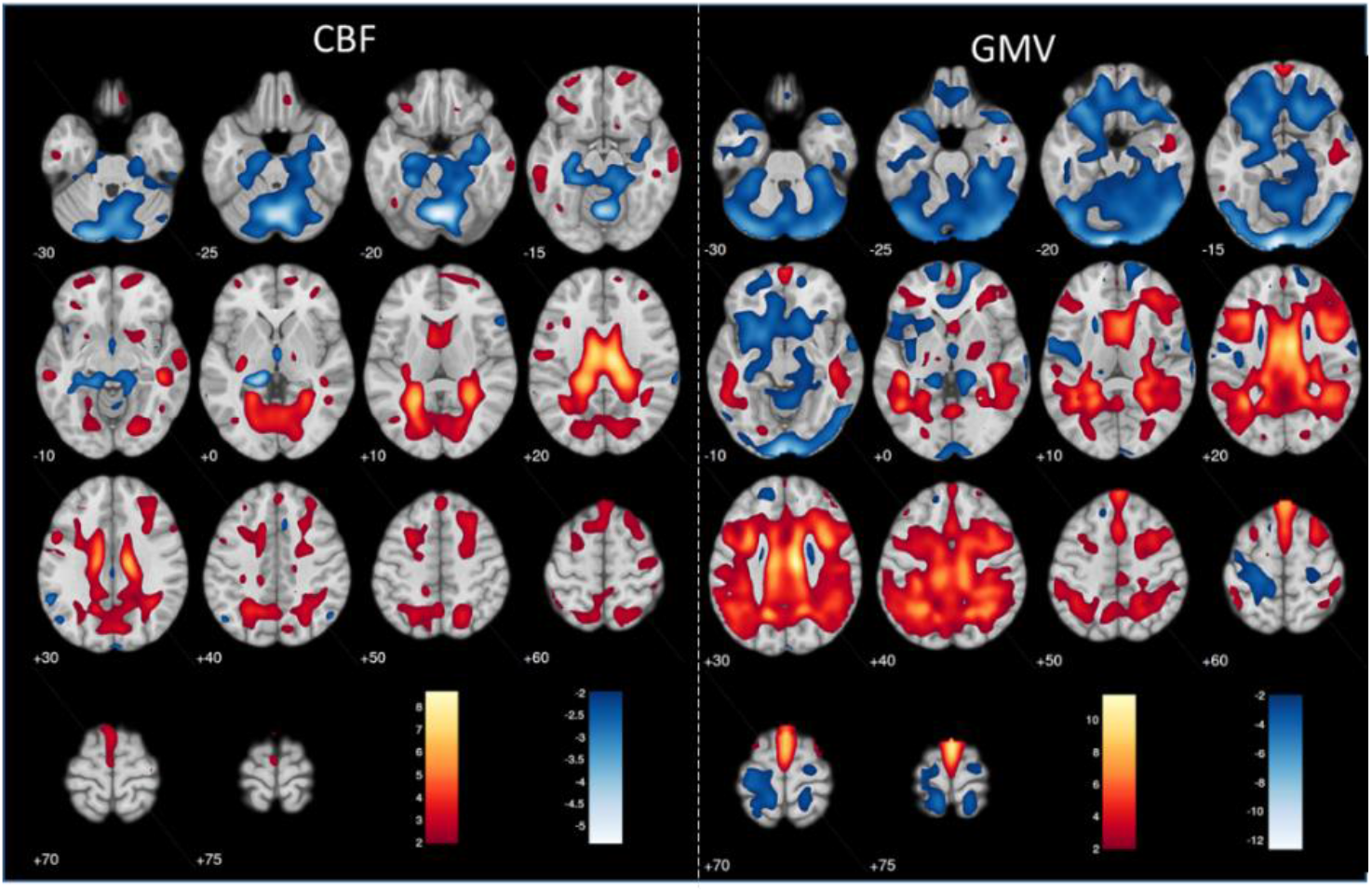
Voxel-wise associations between RSFA and CBF (left panel, Model II) and GMV (right panel, Model V). Maps are thresholded at uncorrected p-values of 0.05 for more complete description of the spatial represnation.

